# The Murphy number: how pitch moment of inertia dictates quadrupedal walking and running energetics

**DOI:** 10.1101/2020.04.24.060509

**Authors:** Delyle T. Polet

## Abstract

Most quadrupedal mammals transition from a four-beat walk to a two-beat run (e.g. trot), but some transition to a four-beat run (e.g. amble). Recent analysis shows that a two-beat run minimizes work only for animals with a small pitch moment of inertia (MOI), though empirical MOI were not reported. It also remains unclear whether MOI affects gait energetics at slow speeds. Here I show that a particular normalization of the pitch moment of inertia (the Murphy number) has opposite effects on walking and running energetics. During walking, simultaneous fore and hindlimb contacts dampen pitching energy, favouring a four-beat gait that can distribute expensive transfer of support. However, the required pitching of a four-beat walk becomes more expensive as Murphy number increases. Using trajectory optimization of a simple model, I show that both the walking and slow running strategies used by dogs, horses, giraffes and elephants can be explained by work optimization under their specific Murphy numbers. Rotational dynamics have been largely ignored as a determining factor in quadrupedal locomotion, but appear to be a central factor in gait selection.

## 1 Background

Despite their incredible morphological diversity, cursorial quadrupedal mammals typically use stereotyped gaits. As speed increases, mammals commonly transition from a four-beat walk at slow speeds to a two-beat trot or pace (where beats are distinct contact events). We see the 4 → 2 pattern across disparate families, such as equids (horses [1]), canids (dogs [2]), bovids (sheep and gazelle [2, 3, 4]), camelids (dromedaries [5]) and antilocaprids [6].

This pattern is surprising from an energetic perspective. A simple accounting of energetic losses in gait is to consider leg contacts as collisions acting on the center of mass (COM) [7]. This perspective explains many phenomena in locomotion, including the pre-heelstrike pushoff in bipedal walking [8], the smooth trajectory of gibbon brachiation [9], why individuals use a flatter running gait in reduced gravity [10], and the leg sequence in transverse galloping [7].

The point-mass collisional perspective posits that frequent, evenly-spaced collisions are better than infrequent, irregular collisions. To optimize work, a quadruped should use as many contacts as possible during a stride; a pronk costs twice as much as a trot, which costs twice as much as a four-beat amble (see supplemental information for a simple derivation).

Why, then, do so many mammals trot? It is unlikely that a slow, four-beat running mode is physically impossible for trotters. The “gaited” horses have been bred to exhibit such gaits^1^. Notable examples are the tölt of the Icelandic horse [11], the amble of the American Saddlebred Horse, and the running walk of the Tennessee Walking Horse [12]. Given the few morphological differences between gaited and non-gaited breeds, it seems less likely that natural populations are physically constrained from performing a four-beat run, and more likely that they reject it (whether through behavioural, developmental or evolutionary programming; e.g. [13]).

In a recent article, Usherwood resolved the paradox by considering the energy of pitching the body [14]. Assuming ground-contact forces are axial to the leg, then foot contact in a four-beat gait induces pitching, but a two-beat gait can avoid it. The question, then, is when do the energetics of pitching outweigh the energetics of COM translation? When pitching energetics dominate, trotting should be optimal, and when translation dominates, tölting should be.

Usherwood [14] showed that the ratio of translational to rotational kinetic energy is related to the dimensionless group

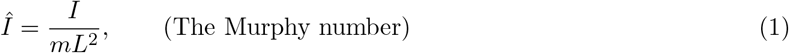

originally defined by Murphy [15] (cited in [16]) with relation to the stability of bounding. In this equation, *m* is body mass, *I* body pitch moment of inertia (MOI) about the COM, and *L* is half the shoulder-hip distance. This dimensionless MOI (called hereafter the “Murphy number” for expediency and in honour of its discoverer), is exactly the ratio of the change in translational to rotational kinetic energy imparted to a free object by a generating impulse perpendicular to L (supplemental information). For *Î* < 1 more rotational energy is imparted than translational, and the opposite is true for *Î* > 1 (figure 1).

**Figure 1:**
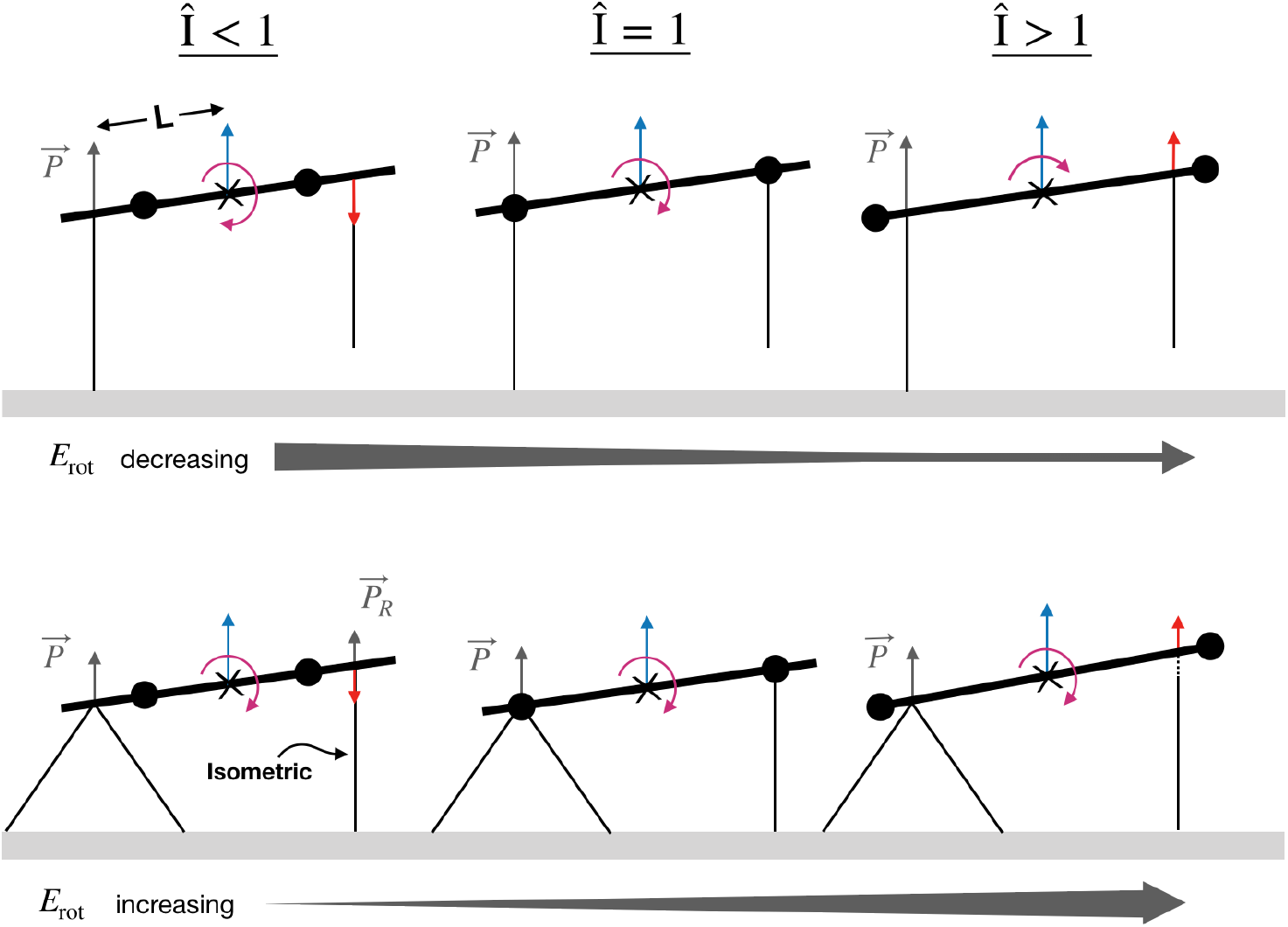
Heuristics showing why Murphy number has opposite effects on walking and running energetics. An impulse 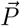 is generated at the hindlimbs and produces an equal change in center of mass velocity (blue arrow) and translational kinetic energy (E_trans_) across all cases. (*Upper row*) In four-beat running as Murphy number increases (left to right), the angular velocity (purple arrow) and rotational energy (E_rot_) decrease. When *Î* < 1, *E*_rot_ > *E*_trans_ and a two-beat gait should be favoured. (*Bottom row*) In four-beat walking, the impulse from hindlimb transfer of support generates a reaction impulse at the forelimbs 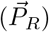 for *Î* < 1, as in this condition the induced forelimb velocity change is downward (red arrow). Because of this, the angular velocity in a four-beat walk does not change as Murphy number increases. However, the rotational energy increases proportionally with MOI. A two-beat gait should be favoured for some *Î* > 1 when *E*_rot_ > *E*_trans_. For *Î* > 1, the impulse 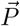 causes a positive change in forelimb vertical velocity. However, if forces are not instantaneous, the forelimb can compensate by reducing its applied force, maintaining a constant pitch rate.

For short stride times, töolting work is related to trotting work by (supplemental information)

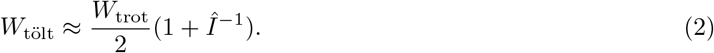

For large Murphy numbers, the point-mass analysis is justified; no energy goes into pitching, and a tölt is cheaper. For very small Murphy numbers, the rotational term dominates, all the energy goes into pitching, and a trot is cheaper. But when *Î* = 1, the cost of tölting and trotting are equal. In general, a four-beat run is optimal when *Î* > 1, but when *Î* < 1, a two-beat run is optimal.

This insight might point to why some mammals deviate from a two-beat run at moderate speeds. Elephants and many primates use a four-beat amble at trotting speeds [17, 18]; giraffes and ring-tailed lemurs transition directly from a walk to a canter [6, 19]. *Î* > 1 implies either that a significant portion of an organism’s mass lies outside its torso, or some mass is positioned a large distance away from the COM (relative to hip-shoulder length). It seems plausible that the large heads of elephants, the long and/or massive tails of some primates, and the long necks of giraffes might push their Murphy numbers beyond unity, but this was not tested by Usherwood [14].

While rotational dynamics and the Murphy number would seem to rectify the two-beat running paradox, it raises another question: why is quadrupedal walking typically four-beat? A mammal using a four-beat walk exhibits pitching of the back [20, 21]. If these rotational energies are large, shouldn’t the same arguments for the trotting-töolting tradeoff apply?

Four-beat walking benefits from distributed contacts interspersed with passive vaulting phases, where the system dynamics in stance resemble a four-bar linkage [22]. To maintain passive vaulting, a pitching torso is necessary, and the pitching direction must be reversed on each transfer of support. This means angular momentum must be absorbed and resupplied with every step^2^. The orientation of the body at transfer of support is predetermined by the geometry of the four-bar linkage, which is independent of the body’s mass or MOI. Likewise, if step-length and speed are predetermined, then the time between hind and fore transfers of support is independent of MOI. Since the rotational speed is independent of MOI, pitching energy should be *proportional* to MOI-not inversely proportional, as in running (figure 1).

We would therefore expect the Murphy number to have the *opposite* effect on the energetics of walking as compared to running. At large *Î*, a two-beat walk should be favoured to avoid costly pitching at the expense of larger COM collisions. At low *Î*, the optimal strategy should be to distribute contacts in a four-beat walk, but switch to a pitch-free two-beat run at higher speeds-the common 4 → 2 pattern.

However, mammals that avoid two-beat running typically do *not* avoid four-beat walking; the walking gaits of elephants, giraffes, and ambling primates appear to be four-beat [17, 23, 18, 24]. It is possible that their gait transition patterns are explained by subtle dynamical effects overlooked by these heuristic arguments.

I examine the energetics consequences of changing Murphy number and speed through trajectory opti-mization of a simple quadrupedal model with a work-based cost function. I also use published data to test the hypothesis that Murphy number is a predictor of differences gait choice between quadrupedal mammals. Finally, using the results of the model, I point to interesting consequences of Murphy number on optimal ground reaction forces, and why point-mass dynamics are insufficient to explain quadrupedal walking.

## 2 Methods

### 2.1 Computational modelling and optimization

The quadrupedal model is planar and based on the methodology of Polet and Bertram [25]. Legs are massless prismatic actuators; limbs cannot generate torque about their respective attachment points to the torso. For simplicity, limb lengths are equal to inter-limb spacing (2*L*), and the COM is located halfway between the fore and hind limbs.

There are a few noticeable differences between the present simulation methodology and that of [25]. First, I constrain the analysis to symmetrical gaits. I compute optimal gaits using trajectory optimization (direct collocation) over the *half* stride cycle. A full stride cycle can be generated by repeating the solution in the second half of the cycle. Since the body is fully symmetrical about the torso center, torso pitch angle can be as high or low as ±π, and I do not impose a limb excursion constraint. Some bipedal solutions emerged as locally optimal on occasion. These were eliminated from the analysis *post hoc*.

Following [25], the present model uses an objective combining limb work with a penalty proportional to the integral of force-rate squared and complementarity violation terms. The force-rate penalty smoothes otherwise impulsive work-minimizing solutions, and is otherwise presently not of interest. Its normalized penalty coefficient is 3 × 10^−5^, 100 times smaller than the value used in [25].

Like the model of [25], the present trajectory optimization setup uses complementarity constraints to allow the optimizer to determine the stepping sequence. Optimizations were carried out with *hp*-adaptive quadrature in GPOPS-II (v. 2.3) and the NLP solver SNOPT (v. 7.5).

I define a non-dimensional stride length and mean horizontal speed as *D*′ = *D/L_H_* and 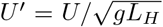 respectively, where *L_H_* is hindlimb length (equal to 2*L* in the model). These correspond to a common normalization seen in the literature [26]. I use the prime superscript (·)′ and the hat diacritic 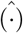 to denote variables normalized by hindlimb length or half inter-limb spacing respectively.

*D*′ was determined from *U*′ through an empirical relationship for walking cursorial mammals [26]:

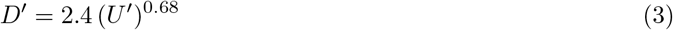

Grid points were selected between 0.25 ≤ *Î* ≤ 10 and 1.5 ≤ *T*′ ≤ 4. The latter represents the lowest and highest stride times observed by [26] among cursorial mammals. *T*′ = *D*′/*U*′ was used as target inputs to the model, as it determines running energetics more directly than speed (see analytical analysis in supplemental information).

An initial search took place at grid points on *T*′ = 0.25 intervals, and *Î* = 0.25, 0.5, 0.75, 1, 1.25, 2, 5, and 10. Afterwards, grid points were added close to identified transition zones between gaits. 50 initial guess were used for each *T*′ > 2.5 condition, while 100 initial guesses were used for *T*′ ≤ 2.5. Convergence was difficult at the slowest speeds (*T*′ = 4), and several outliers were identified as isolated gaits of a certain number of beats surrounded by solutions with a different number of beats. To these solutions, another 50 guesses were added to better converge to the optimal solution. Initial guesses were formed by selecting from a uniform random distribution across each variable’s range at 16 uniformly-spaced grid points.

For a given parameter combination, the lowest-cost solutions were selected among all local minima discovered. The beat number was determined *post hoc* by looking at peak negative power during the stride. Defining a beat as peak negative power is consistent with the collisional gait perspective, which points to mechanisms of energy loss and approximates them as impulsive events [7, 27]. Setting normalized (negative) power as

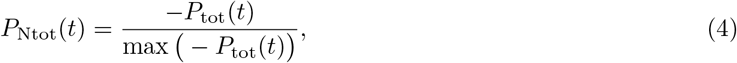

where *P*_tot_ is instantaneous net power from all actuators, the number of beats were the number of local maxima in *P*_Ntot_(*t*) > 0.3. If two maxima were less than 0.03 *T* apart, the greater maximum among them was counted as a single beat. This method eliminated some noise while selecting only the largest events of energy loss as a “beat”. However, it is somewhat arbitrary and the shape of gait “zones” changes to some extent depending on tolerances (see supplemental information for results using other tolerances).

Limb contact for a given limb was defined as its GRF > 0.01 *mg*. Walking was defined as having duty factor > 0.50 in at least one pair of limbs (fore or hind), with running being all other cases. Although this distinction aligns with Hildebrand’s use of the terms “walk” and “run” for symmetrical gaits [12], there are examples in nature of “grounded running” where the COM bounces as in a run, but duty factors exceed 0.5 [28, 17].

### 2.2 Calculations of empirical moments of inertia

The relevant pitch moments of inertia (MOI) about the COM were taken during standing, and were derived from values reported in the literature (Supplementary table S1). Alexander [29] measured whole body MOI for an Alsatian dog directly and reported the normalized value along with body mass. The reference length (not reported by the author) was derived from figure 12 in [2], which Alexander used as a reference [29]. Whole body MOI for the Dutch Warmblood was calculated from [30] using figure 1 from that study as a guide for limb and head orientation.

For the elephant and giraffe, no direct MOI measurements are available, but some studies report estimates using 3D models. [17] provide measured masses and estimated moments of inertia for elephants. The shoulder-hip length was calculated by scaling their reported limb lengths to figure 1 in [31]. Estimated MOI was also derived for a horse and giraffe from [32]. Shoulder and hip locations were estimated by comparing their figure 1 to skeletal drawings or mounts. COM position was assumed to lie along the shoulder-hip line, and its bias towards the forelimbs (*m*′_*F*_ in [25]) was determined from calculations using [30] for the horse (*m*′_*F*_ = 0.50) and ground reaction forces from [23] for the giraffe (*m*′_*F*_ = 0.65). The horse MOI was used to ground-truth the estimate from [32], and yielded *Î* = 0.80, similar to the empirical value of 0.82 (supplementary table S1).

## 3 Results and Discussion

Figure 2a shows optimal gaits at parameter combinations of *Î* and *U*′. Optimal gaits generally fall into four large regions. At high *Î*, four-beat runs and two-beat walks are optimal. At low *Î*, the reverse is true. While the cutoff between two- and four-beat runs is approximately *Î* ~ 1, as predicted by equation (2), the transition *Î* for four-beat to two-beat walking increases from about *Î* = 1 at the highest walking speeds to *Î* = 2 at the slowest speeds examined.

**Figure 2.**
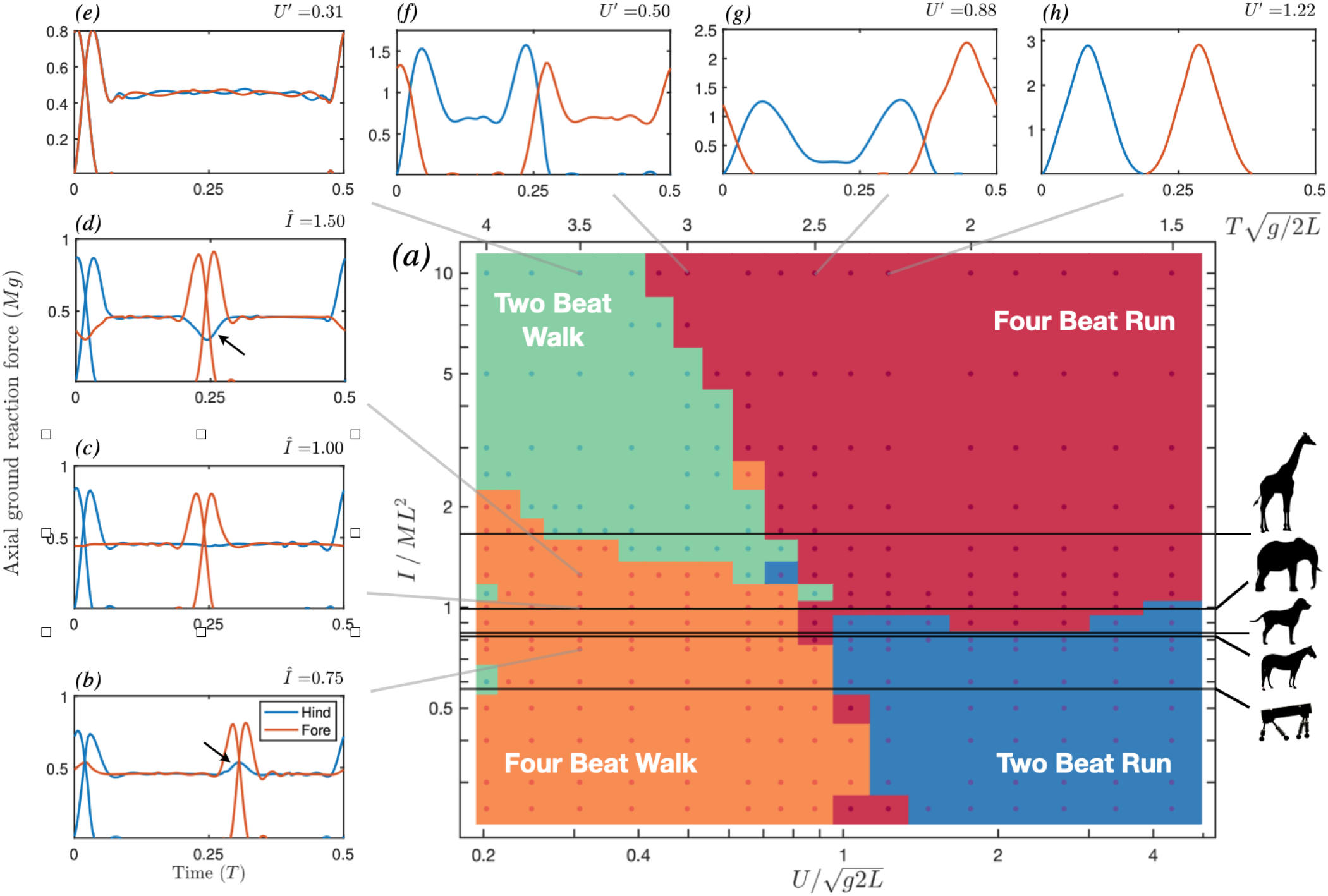
Optimal gaits for parameter combinations of *Î* and *U*′. (a) Gaits largely fall into four main regions. At high *Î*, four-beat runs and two-beat walks are optimal. At low *Î*, the reverse is true. Dogs and horses have *Î* < 1, and exhibit a four-beat walk and two-beat run, as predicted. Likewise, a robot with low *Î* finds 4 → 2 beat transition to be energetically optimal [33]. Elephants, with *Î* = 1, exhibit a four-beat gait regardless of whether walking or running [17]. Giraffes have the most extreme *Î* examined here and do not use a two-beat run [34]. Despite their large *Î*, a four-beat walk remains optimal at slow speeds. (*b-h*) Limb-axial ground reaction forces (GRF) for a number of optimal solutions (animated in supplemental videos 1 and 2). A half-cycle is shown in each case; the solution is repeated during the second half-cycle, but mirrored in the sagittal plane. (*b-d*) Transfer of support in one pair of limbs during four-beat walking induces a vertical reaction force at the other pair. (*b*) When *Î* < 1, the vaulting limb exhibits an increase in GRF (arrows) to cancel the negative reaction force and maintain its length. (c) At *Î* = 1, the vaulting limb does not exhibit a change in force. (*d*) At *Î* > 1, the vaulting limb sees a reduction in midstance force. (e) At slow speeds and high *Î*, a two-beat walk is optimal. (f) As speed increases with *Î* = 10, a four-beat “running” solution emerges, with single limb vaulting swapping between fore and hind limbs. (*g*) At higher speeds, the optimal gait is a hybrid between vaulting in hind and bouncing in front, reminiscent of the slow tölt [11]. (h) At still higher speeds, a typical fast tölt pattern emerges.

These findings support the hypothesis that Murphy number has opposite effects on the optimality of gait in walking and running. In both walking and running, there is a tradeoff between distributing collisions between multiple contacts (favouring four-beat gaits) and avoiding work to pitch the body (favouring two-beat gaits). During running, the possible energetic losses from pitching increase as Murphy number *decreases*, while in walking, these losses increase as Murphy number *increases*.

For dogs and horses (*Î* ~ 0.8), the tradeoff means it is optimal to use a four-beat walk and two-beat run, as these animals typically do. Indeed, even a quadrupedal robot with a small Murphy number finds the same 4 → 2 beat transition to be optimal [33]. For elephants, which have *Î* ~ 1, the tradeoff seems to favour a four-beat gait regardless of speed. In reality, these animals only use four-beat gaits, and it is difficult to distinguish their transition from walking to running [17].

A four-beat run is predicted for dogs and horses right before the transition to two-beat running. The optimal solution is borderline between walking and running: the ground reaction forces exhibit the double-hump profiles characteristic of walking, and the duty factor is slightly less than 0.5. Increasing the force-rate penalty extends the duty factor at the walk-trot transition comfortably into the walking range, and better matches empirical data [25], highlighting that minimizing limb-work alone does not fully explain gait choice.

The giraffe has the most extreme Murphy number by far of all the mammals investigated here. It also has unusual gait patterns, exhibiting only two gaits, the walk and the gallop^3^, with no intermediate trot [34, 6, 4]. However, figure 2a predicts that giraffes should have *three* distinct gaits: a four-beat walk at slow speeds (*U*′ < 0.34), a two-beat walk at higher speeds (0.3 < *U*′ < 0.7) and a four-beat run at higher speeds (0.8 < *U*′). The walk-run transition point appears sensible, as Basu *et al.* [35] report a walk-gallop transition speed^4^ of *U*′ ≈ 0.8.

Walking giraffes exhibit a mean hind-fore phase offset of 0.14 (range 0.09-0.2) [21, 23], above the 0.0625 limit for a pace given by Hildebrand [12]. These observations of a four-beat gait are for an extremely slow normalized speed (0.14 < *U*′ < 0.30), matching the region where the work-minimizing model predicts a four-beat walk (figure 2a). Is there any evidence of giraffes using a two-beat walk at intermediate speeds?

The walk of the giraffe has been described in two-beat terms, including “rack-like” [34] or as a pace [36]. However, without quantifying the phase relationship or speed at which these observations were made, it is not clear whether these represent the same gait quantified as four-beat in other studies [23, 21]. It seems difficult to elicit walking speeds above *U*′ = 0.3 for giraffes in captivity [37, 23]. Indeed, Innis [34] reports that wild adult giraffes seem to use only two modes; a “leisurely” walk or a fast run. This leaves a large gap (0.3 < *U*′ < 0.8) where giraffe gait has not been quantified, approximately where figure *2a* anticipates a transition to a two-beat walk.

We should not place too much weight on the model’s exact quantitative predictions in this case. Giraffes occupy a region of *Î*-*U*′ space where subtle changes in MOI can have profound changes on 4 → 2 walk transition speed. Added to the fact that *Î* is highly sensitive to L (a 5% change in L can lead to a 10% change in *Î*), the predicted four-beat to two-beat walk transition could vary substantially with measurement error. The walk-run transition speed, however, is less sensitive to choice of *Î*.

While the model correctly predicts the absence of a two-beat run, giraffes do not use the predicted symmetrical four-beat run either, instead opting for a three-beat canter or four-beat gallop. While the present symmetrically-constrained model could not reproduce these asymmetrical gaits, a collision-based analysis predicts that a canter should be optimal at intermediate running speeds for a long-limbed animal with a high Murphy number, such as a giraffe [14].

### 3.1 Changes in walking strategy with Murphy number and speed

An interesting effect in walking can be observed as Murphy number increases. During transfer of support on one set of limbs (*e.g.* the hind pair), the leg at the opposite end of the body (*e.g.* forelimbs) will exhibit increased reaction force if *Î* < 1 and decreased reaction force if *Î* > 1 (figure 2b-d). Why does this occur?

An impulse at the hindlimbs simultaneously causes the body to translate upwards and pitch down. Depending on how much the impulse causes rotation *vs* translation, the net effect at the instant of the impulse may be to push the forelimbs up or down. For *Î* < 1, an impulse at the hips cause the shoulders to *descend*; at *Î* > 1, the same impulse causes the shoulders to *ascend*; and at *Î* = 1 the shoulders remain (momentarily) stationary (figure 1).

During walking, it is advantageous for the vaulting limb to maintain its length; a change in length while providing axial force implies that the limb is performing work. The strategy, then, is for the vaulting limb to cancel the force it feels from the limbs undergoing transfer of support. For *Î* < 1, double-stance contact induces a downward force on the vaulting limb, which can respond by increasing its applied force so as to maintain its length and perform no work (figure 2*b*).

For *Î* > 1, double-stance induces an upward force on the vaulting limb. The vaulting limb therefore responds by *reducing* its applied force, maintaining its length (figure 2*d*). At some large *Î*, this strategy will fail; the vertical joint reaction force induced on the vaulting limb exceeds its upward ground reaction force (~ 0.5*mg*). This constraint does not seem to govern the transition to two-beat walking, however. *Î* = 1.5 and *U*′ = 0.3 is at the border between four-beat and two-beat walking (figure 2a), yet the vaulting limb only reduces its applied force by 0.2*mg* (figure 2d).

Two-beat walking forms a wedge in the upper-left corner of figure 2*a* (example sequence in figure 2e), and the transition to four-beat running occurs at lower speeds as the Murphy number increases. The four-beat “running” solution at *U*′ = 0.5, *Î* = 10 demonstrates why (figure 2*f*; see also supplemental video 2). The solution is to perform a single-limb vault over forelimbs, then hindlimbs, and repeat this pattern. This solution is feasible because the Murphy number is so extreme that the body barely pitches during single stance, even though it is supported only at one end.

As we increase Murphy number, we approach the limit where any pitching can be effectively ignored. In this limit, we expect all gaits to be four-beat; it is analogous to a point mass biped with half the stride length. At slow speeds, a point mass biped should use a vaulting walk to minimize work [38]. With an extra set of legs, it can reduce contact losses by taking twice as many steps per stride (similar to the solution observed in figure 2*f*). At intermediate speeds, a point-mass biped should use a hybrid gait: a pendular run with single-leg contacts [38]; again, adding two legs means we simply halve the stride length. The simulation discovers a similar hybrid gait (figure *2g*)-reminiscent of the slow tölt [11]. The same logic applies to impulsive running, the minimal-work high-speed gait for a point-mass biped, resulting in a familiar fast tölt (figure 2*h*). The extreme case of *Î* = 10 has sufficient pitching energies that a two-beat walking gait is optimal at slow speeds (figure 2*e*); as we further increase *Î*, we expect the four-beat transition speed to decrease.

## 4 Summary and Conclusions

Contact forces axial to the a quadruped’s legs pitch its body, unless compensated by a counter torque. The Murphy number parameterizes the tendency of these contacts to pitch the body *vs* accelerate the center of mass. Large Murphy numbers result in less energy going into pitching *versus* translation by single limb contact. As a result, four-beat gaits are favoured, which reduce the collisional cost of changing COM momentum. At lower Murphy numbers, the opposite is true, and more oscillation of the COM is worth the price for avoiding costly pitching. The transition point between two-beat and four-beat running occurs close to *Î* = 1, matching an analysis by Usherwood [14].

However, Murphy number has the opposite effect on walking energetics, due to the geometric constraints of four-beat walking and the ability of the vaulting limb to counteract some of the effects felt by transfer of support at the opposite pair of limbs. Altogether, the work-based model correctly predicts the walking and slow running gaits selected by dogs, non-gaited horses, elephants, and a quadrupedal robot. It also correctly predicts that giraffes should use a slow four-beat gait and avoid trotting at high speeds. It does not (nor can it) predict a canter as the slow-running gait of choice for giraffes, and predicts that giraffes should use a two-beat walking gait at intermediate to fast walking speeds (for which there is currently no data).

Point-mass collisional dynamics predict that all quadrupedal gaits should be four-beat with alternating single stance contact. It is only by considering pitching dynamics that other gaits emerge as energetically optimal solutions. Except for some specialized gaits-trotting, cantering, and possibly transverse gallopings [7]–, the net torque about the COM is appreciable and energetically costly. Furthermore, these non-pitching gaits may be commonly used *precisely because* pitching would otherwise be extremely costly. Pitching may be so important energetically, that the optimal solution is often to render it absent.

## Data Availability

The dataset supporting this article has been uploaded as part of the supplementary material (Table 1), and on Zenodo at https://doi.org/10.5281/zenodo.3765877.

## Acknowledgements

I would like to thank Jim Usherwood for inspirational discussions on this topic, John Bertram and Ryan Schroeder for helpful comments on early drafts, and Jessica Theodor for providing computational resources.

## Footnotes

1 Four-beat gaits are desirable as they exhibit less oscillation of the COM, resulting in a smoother ride for the human in the saddle.

2 It is possible that the braking impulse freely transfers some of the rotational energy into translation, though for simplicity this is assumed small.

3 There is some debate whether the canter, also used by giraffes, is a distinct gait or merely a slow gallop.

4 As *U*′ ≈ 0.8 was the slowest running gait observed, and the authors did not report on walking gaits in that study, it is unclear if this is truly the transition speed.

