## Supplemental Information for "The Murphy number: how pitch moment of inertia dictates quadrupedal walking and running energetics"

#### 1 The relationship between the Murphy number and kinetic energy

Assume that an impulsive force, applied at $t = 0$ at a distance $L$ from the center of mass (COM), generates a torque about the COM. We call the component of the force perpendicular to the moment arm $F$. This component causes a change in the translational and rotational velocities of

$$F = m\frac{\mathrm{d}V}{\mathrm{d}t} \tag{S1}$$

$$FL = I\frac{\mathrm{d}\omega}{\mathrm{d}t} \tag{S2}$$

where $V$ is the center of mass velocity and $\omega$ is the angular velocity about the center of mass. Equating $F$ in these equations and separating differentials yields

$$\int_{t=0-}^{t=0+} m\,\mathrm{d}V_c = \int_{0-}^{0+} F\,\mathrm{d}t = \int_{t=0-}^{t=0+} I/L\,\mathrm{d}\omega \tag{S3}$$

Assuming $V(0^-) = \omega(0^-) = 0$, and setting $V(0^+) = V$ and $\omega(0^+) = \omega$ as unknowns, we find

$$V = I\omega/(mL). \tag{S4}$$

We would like to know the ratio between the translational and rotational kinetic energy imparted to the body. This is

$$\frac{K_t}{K_r} = \frac{mV^2}{I\omega^2} \tag{S5}$$

$$= \frac{mI^2\omega^2}{Im^2L^2\omega^2} \tag{S6}$$

$$= \frac{I}{mL^2} \equiv \hat{I} \tag{S7}$$

Therefore, the Murphy number is exactly the ratio of the translational to rotational kinetic energy imparted by the component of the impulsive force perpendicular to the moment arm.

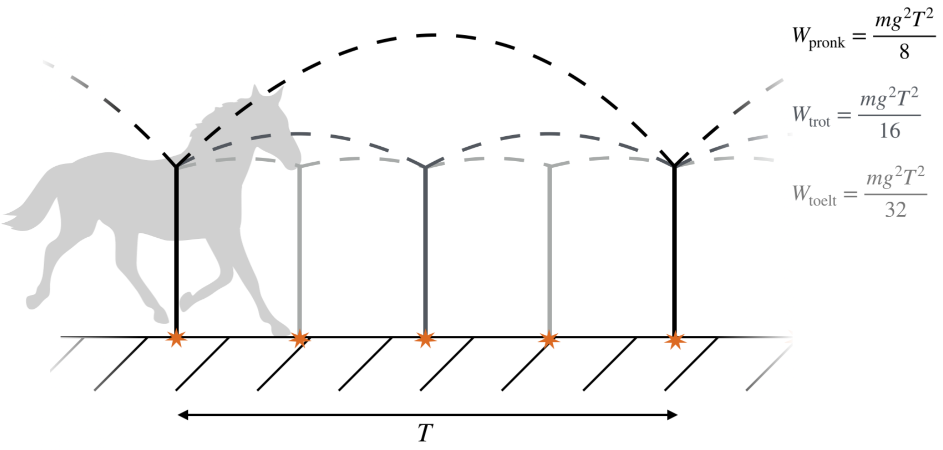


Figure S1: Comparison of work done for three gaits with the same stride period $T$ but a different number of contact sequences. Compared to a one-beat pronk, a two-beat trot can cut work in half, while a four-beat tölt can cut work down to a quarter of the cost, if only translational energy of the center of mass is considered in a collisional point-mass model similar to [2].

#### 2 Analytical derivation of four-beat *vs* two-beat running work

The recent article by Usherwood [1] gives considerable insight into the energetics of quadrupedal running. It relied on an assumption that the body remains horizontal; in other words, that time between contacts is infinitesimally short. However, the rotation imparted by an impulse at the hindlimbs can perhaps be mitigated by making contact when the body is at an angle to the horizontal. In this way, the necessary vertical impulse is generated to redirect the center of mass from down to up, but less energy is imparted to rotation than may be necessary. Do Usherwood's claims hold when finite pitching is considered?

First, we assume that at any given contact, all kinetic energy due to vertical translation is lost immediately and passively[1], and that the actuation is impulsive and perfectly pseudoelastic (that is, it resupplies exactly the lost energy back into the system, following [2]) The energetic cost is exactly the resupplied energy, or the positive work of the actuators. We also assume that all actuation is vertical; this is consistent with work-minimizing running gaits on point-mass systems, where any fore-aft actuation needlessly decelerates and then reaccelerates the center of mass [2]. Consequently, the horizontal velocity is immaterial, and the only kinematic parameter which we must provide is the stride period $T$.

When an animal is modelled as a simple point mass in running with inelastic strut-like legs (figure S1), the vertical landing speed of the center of mass ($V$) is lost at contact. Lost energy must be resupplied by positive work, at a cost proportional to $mV^2/2$ with every foot contact[2]. How many contacts should a quadruped use? The time between contacts is given by ballistics as $T_c = 2V/g$. Given a stride time $T$, the touchdown speed then depends on the number of contacts $n$ during a stride as $nT_c = T$ (assuming infinitesimal contacts). The touchdown speed is thus $V = gT/(2n)$; we can therefore sum across all contacts

[1] This is **not** necessarily true in the case of a distributed mass, as it would be for a point mass during impulsive running with vertical legs. [2] Elastic storage reduces this cost, but higher kinetic energy still usually means more energy lost at contact.

to get the positive work required in a stride:

$$W_{\text{stride}} = n\frac{m}{2}\left(\frac{gT}{2n}\right)^2 \tag{S8}$$

$$= \frac{mg^2T^2}{8n} \tag{S9}$$

Since total positive work in the stride is inversely proportional to $n$, the quadruped should use as many contacts as possible. What is the relative cost of trotting *vs* tölting[3]? From equation (S9), we can state

$$W_{\text{trot}} = \frac{m(gT)^2}{16}, \tag{S10}$$

$$W_{\text{tölt}} = \frac{m(gT)^2}{32} = W_{\text{trot}}/2 \tag{S11}$$

This is the central paradox from the point-mass collisional perspective. Because of the additional contacts, a four-beat gait should always be cheaper than a two-beat gait, but the latter is often preferred.

This analysis has so far ignored pitching energetics. Trotting can avoid pitching by providing equal and opposite torques about the center of mass at contact. But in tölting, pitching is unavoidable (assuming no torque is transmitted to the body by the limb about its attachment point).

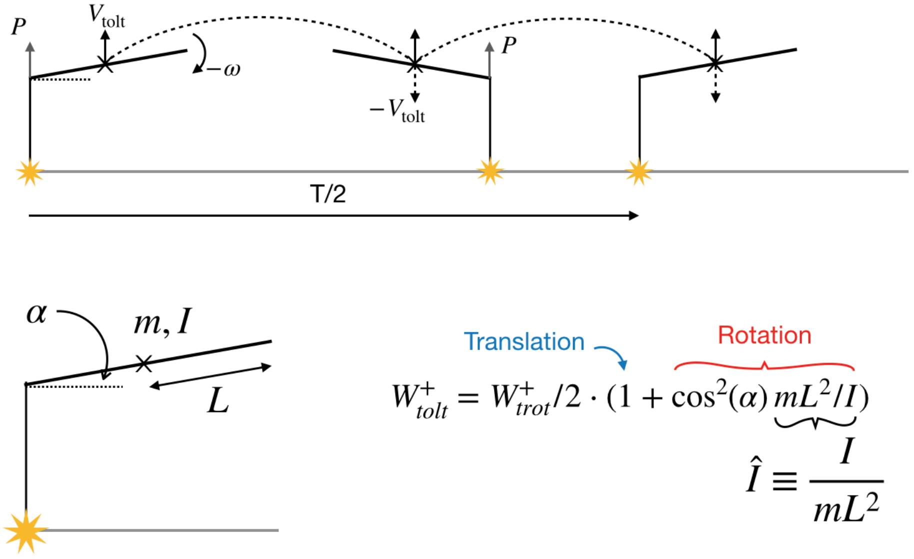


Figure S2: **A collisional model of tölting that includes pitching.** Vertical contacts generate impulse $P$ that induces the body to pitch at angular speed $\omega$. The model is defined with contact pitch angle $\alpha$, body mass $m$, pitch moment of inertia $I$ and half-support spacing $L$. An analysis of the energetics yields two terms; one associated with translational kinetic energy, and related to figure S1, and another associated with rotational kinetic energy in which the Murphy number ($\hat{I}$) appears.

We want to know the angular velocity of pitching $\omega$ due to the impulsive limb contact. We give our quadruped a pitch moment of inertia of $I$ and an even mass distribution between the fore and hind limbs, with a hind-to-fore distance of $2L$. Contact occurs at a pitch angle $\alpha$ and, due to the symmetry of our

[3] In this section I use trotting and tölting to refer to generalized symmetrical two- and four-beat running gaits respectively.

organism, we can set an equal and opposite contact angle for the next limb contact (figure S2). Therefore, the time between contacts must satisfy

$$\frac{T}{4} = \frac{2\alpha}{\omega} = \frac{2V_{\text{tölt}}}{g} \tag{S12}$$

With the introduction of the unknown $\alpha$, we must introduce one more equation to solve for $\alpha$ and $\omega$. The impulse equations provide this. A vertical impulse $P$ at the forelimbs creates a change in linear momentum of $P = m(V^+ - V^-)$, where $^-$ and $^+$ denote a variable before and after contact, respectively. With the assumption that all kinetic energy is lost at contact, we can set $V^- = 0$. We've also defined $V^+ = V_{\text{tölt}}$, and so

$$P = mV_{\text{tölt}}. \tag{S13}$$

This same impulse generates a change in angular momentum. The component perpendicular to the body axis is $P \cos \alpha$, and so the change in angular momentum is $LP \cos \alpha = I(\omega^+ - \omega^-)$. Again, with the assumption that all kinetic energy is lost at contact, and by defining $\omega \equiv \omega^+$, we have

$$LP \cos \alpha = I\omega. \tag{S14}$$

Solving for $P$ in the relations S13 and S14 and equating, we get

$$\omega = \frac{mV_{\text{tölt}} L \cos \alpha}{I}. \tag{S15}$$

We insert S15 into S12 to get a constraint on $\alpha$

$$\frac{\alpha}{\cos \alpha} = \frac{mLT^2 g}{I64}. \tag{S16}$$

If we define two non-dimensional parameters $\hat{I} \equiv I/mL^2$ and $\hat{T} \equiv T\sqrt{g/L}$, then S16 simplifies to

$$\frac{64\alpha}{\cos \alpha} = \frac{\hat{T}^2}{\hat{I}}. \tag{S17}$$

$\hat{T}$ is a normalized stride period and $\hat{I}$ is the Murphy number [1]. equation (S17) tells us that as $\hat{I} \to \infty$, $\alpha \to 0$ for a given $\hat{T}$; in other words, a very large moment of inertia will involve no pitching of the body, as we would expect. For a given $\hat{I}$, a short stride period ($\hat{T} \to 0$) also results in $\alpha \to 0$; in other words, we can use the small angle approximation if we assume $\hat{T}$ is sufficiently short, as we might expect in a fast running gait.

Since four steps are taken, with kinetic energy being lost in each step, the positive work due to translational and kinetic energy is

$$W_{\text{tölt}} = 4(mV^2/2 + I\omega^2/2). \tag{S18}$$

We use equation (S12) to find $\omega$ and equation (S15) for $V_{\text{tölt}}$. Inserting these values into S18, we get

$$W_{\text{tölt}} = \frac{m(gT)^2}{32} + \frac{(mLgT \cos \alpha)^2}{32I}. \tag{S19}$$

The first term is due to translation and depends only on the stride time (given gravity and mass). The second term is due to energy going into rotation, and now depends on stride time *and* the relative values of

$I$, $m$ and $L$. Recognizing the first term as equation (S11), we can simplify to

$$W_{\text{tölt}} = \frac{W_{\text{trot}}}{2}(1 + \hat{I}^{-1}\cos^2\alpha). \tag{S20}$$

Using the small angle approximation (justified from equation (S17) for a short stride period), the equation simplifies to

$$W_{\text{tölt}} \approx \frac{W_{\text{trot}}}{2}(1 + \hat{I}^{-1}), \tag{S21}$$

which is equation (2) in the main text. It tells us that four-beat running at small Murphy numbers involves substantial torques from leg contact, generating large rotational energy that must be absorbed at the next contact. Trotting has the advantage of eliminating pitching completely, but at the cost of fewer leg contacts.

Figure S3 shows $\log_{10}(W_{\text{tölt}}/W_{\text{trot}})$ over a range of stride times and moments of inertia. The changeover at $\hat{I} = 1$ holds for $\hat{T} \lesssim 3$. As $\hat{T}$ continues to increase, tölting remains optimal over trotting for Murphy numbers $< 1$, until about $\hat{T} = 6$, where tölting is always optimal regardless of Murphy number.

For large stride times. costs due to vertical velocity discontinuities become extremely expensive, while the body's rotational velocity is relatively low; both these factors favour tölting. While running at such large stride times is not common (the slowest stride times in walking mammals are at $\hat{T} \approx 6$ [3]), this analysis illustrates that other factors come into play at slow speeds that may make a pitching gait more favourable, even for small $\hat{I}$. As shown in the main text, here a four-beat walk is optimal.

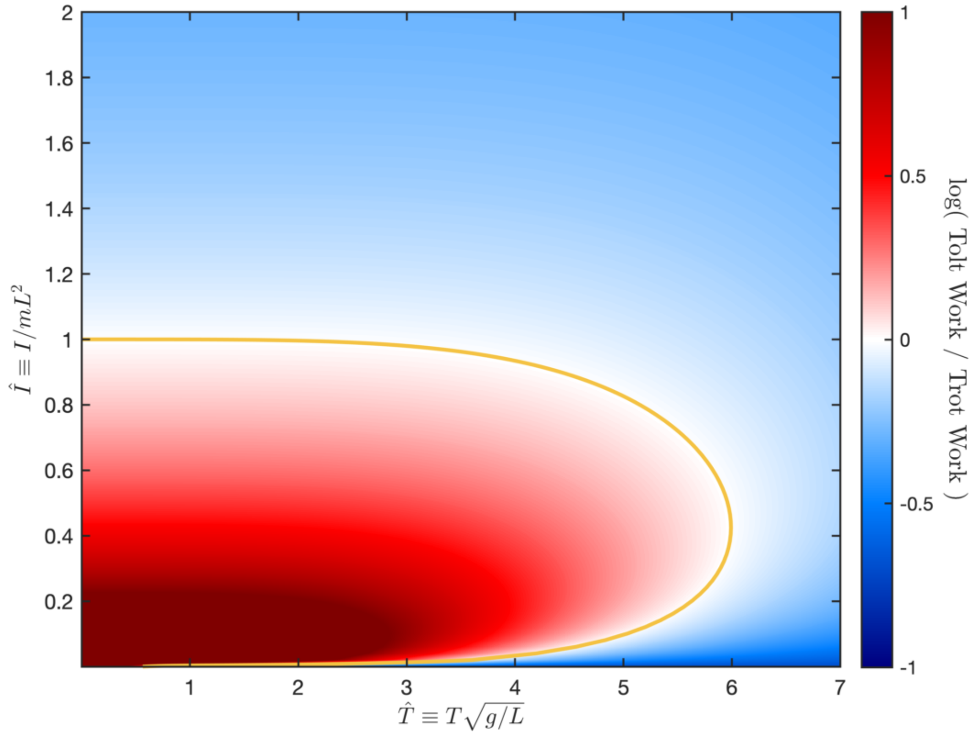


Figure S3: $\log_{10}$ **of the ratio between tölting and trotting costs are compared across a range of combinations of normalized stride time ($\hat{T}$) and Murphy number ($\hat{I}$).** Note here that an increase in stride time (left to right) normally results in a speed *decrease* in organisms. Above 0 (red), trotting is less expensive; below 0 (blue) tolting is less expensive. As stride period increases, tölting is optimal even for $\hat{I}$ slightly less than 1, because costs due to vertical velocity discontinuities become extremely expensive, while the body's rotational velocity is relatively low.

### 3 Table of Empirical Morphological Data

Table 1: Morphological parameters for several mammal species used in this study.

| Species | $I$ (kg m$^2$) | $L$ (m) | $m$ (kg) | $\hat{I}$ | MOI Measurement Method | Primary Source |
|---|---|---|---|---|---|---|
| *Canis lupus domesticus (Altsatian)* | 2.02 | 0.31 | 25 | 0.84 | Direct | [4] |
| *Equus ferus caballus (Thoroughbred)* | 142 | 0.68 | 383 | 0.80 | Indirect (3D model) | [5] |
| *Equus ferus caballus (Dutch Warmblood)* | 229 | 0.73 | 525 | 0.82 | Direct (Segmented) | [6] |
| *Loxodonta africana* | 2005 | 0.84 | 2831 | 0.99 | Indirect (3D model) | [7] |
| *Giraffa camelopardalis* | 1778 | 0.81 | 1611 | 1.66 | Indirect (3D model) | [5] |

### 4 Sensitivity analysis for gait detection tolerances

To detect beats, two tolerances were set: minimum peak height (set at 0.3 times maximum negative power), and minimum distance between peaks (set at 0.03 $T$). Figure S4 shows how different tolerances affect the shape of gait zones. Only slow speeds– especially at low $\hat{I}$– are greatly affected by changes in tolerances. Some solutions near the two-beat to four-beat transition are also affected. At slow speeds, some solutions exhibited multiple peaks in negative work in short succession at transfer of support. With low tolerances, these peaks are counted as unique beats, leading to six- and eight-beat walks. The rest of the parameter space is unaffected by changes in tolerance.

### 5 Interpolation scheme for gait zones

Simulations were time-intensive, so to reduce computational time missing solutions for combinations of $\hat{I}$ and $U'$ were inferred from surrounding solutions. However, standard interpolation schemes generally assume continuous numerical data, whereas the data here is discrete (for example, a 1.5-beat gait has no meaning here, nor does a three-beat cycle for a symmetrical solution). Instead, I used an interpolation scheme based on data points immediately surrounding the missing data. The algorithm was as follows:

1. For a missing point, find its immediate surrounding values
2. If all surrounding values are equal to each other ($x$), or missing, then replace the current missing point with $x$. Otherwise, leave the point as missing and move on to step 1 for the next missing point
3. Once steps 1 and 2 have iterated over the initial set of missing points, find all remaining missing points. For each of the remaining missing points, find its immediate surrounding values
4. If all surrounding values are equal to each other ($x$), with the exception of up to one point, then fill the missing point with $x$.

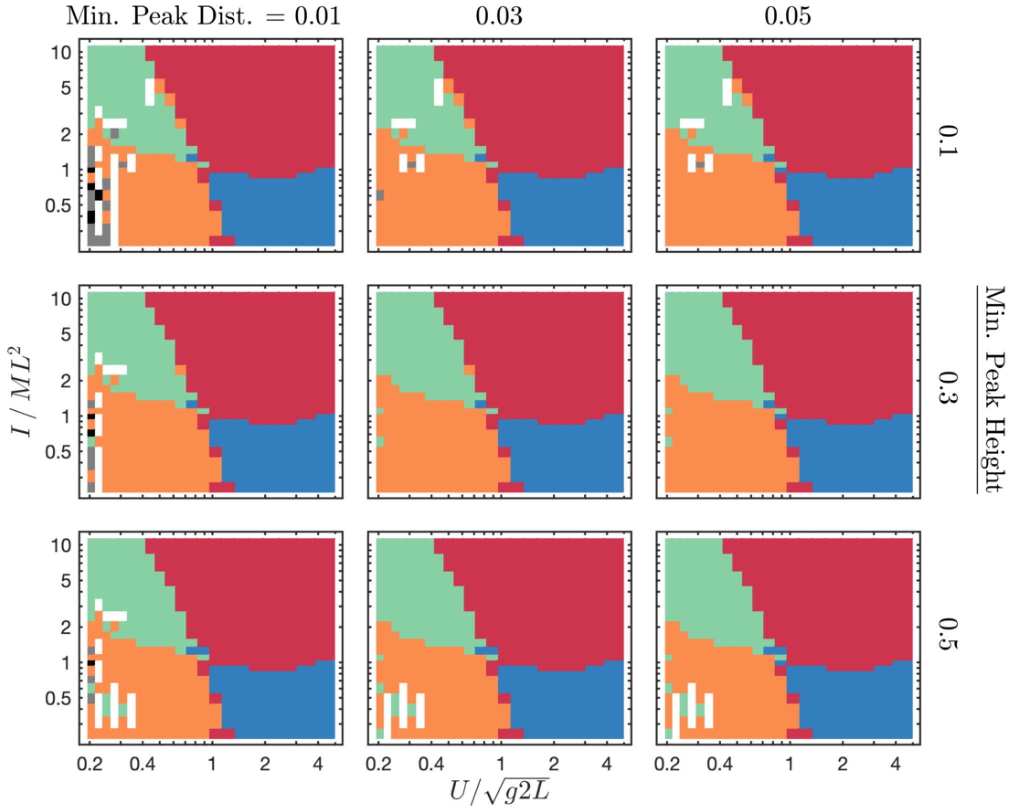


Figure S4: As beat detection tolerances change, the shape of gait zones change at slow speeds and at the two-beat walk to four-beat run transition. Elsewhere, gait zones do not change. Key: green, two-beat walk; red, four-beat run; orange, four-beat walk; blue, two-beat run; grey, six-beat walk; black, eight-beat walk; white, no data, and value could not be interpolated.

This algorithm fills in missing data by matching them to the discrete gaits of their neighbours. All missing data points completely enclosed by a certain gait type would match that gait type, while missing data at gait transition zones would not be filled unless surrounded overwhelmingly by one gait type. I do not claim that this algorithm is a good general-purpose data-filling tool. The reader can judge from figure 2 in the main document whether it filled data satisfactorily for this study.

For Min. Peak Distance = 0.03 and Min Peak Height = 0.3, simulations were completed to fill the entire space at a satisfactory resolution, without any missing points. For the tolerance sensitivity analysis (figure S4), the algorithm could not discern some solutions at gait transitions, and those data points remain blank.

### 6 Descriptions of supplemental videos

Supplemental Video 1: Three walking sequences at $U' = 0.3$ show how four-beat walking does not look qualitatively different as $\hat{I}$ increases. However, the forces of the vaulting limb during transfer of support of the opposite pair do change systematically (figure 2*b-d*). Animations are adjusted so that one stride cycle takes 4 seconds. Leg contact is displayed when ground reaction force exceeds $0.05\,mg$.

Supplemental Video 2: Four sequences at $\hat{I} = 10$ show how gait changes with increasing speed at a large Murphy number. At low speeds, a two-beat gait is optimal. At higher speeds, a gait emerges with DF $< 0.5$ (a run by the classic Hildebrand definition) but with alternating phases of walking-like vaulting between four-and hind limbs. Around $U' = 0.9$, a hybrid gait emerges, with a vaulting phase in hindlimbs and bouncing phase in forelimbs. At still higher speeds, a four-beat run emerges. The ground-reaction forces associated with these cases are shown in figure 2*e-h*. Animations are adjusted so that one stride cycle takes 4 seconds. Leg contact is displayed when ground reaction force exceeds $0.05\,mg$.
